# Queen number shapes worker longevity, fecundity and gene expression in the invasive, highly polygynous ant *Tapinoma magnum*

**DOI:** 10.1101/2024.07.21.604453

**Authors:** Anna Lenhart, Megha Majoe, Sibel Selvi, Thomas J. Colgan, Romain Libbrecht, Susanne Foitzik

## Abstract

In social animals, reproductive activity and ageing patterns are influenced by group composition. A well-documented phenomenon in monogynous (one-queen) insect societies is that queen presence affects worker fecundity and longevity. Little is known about whether and how workers respond to queen number variation in polygynous (multi-queen) species and how their queens age. We created queenless, one-queen, and two-queen colonies of the invasive, polygynous ant *Tapinoma magnum* to examine worker survival, fecundity, oxidative stress resistance, and fat body gene expression. Additionally, we compared fecundity and brain and fat body transcriptomes of young and old queens. Queenless workers experienced the highest mortality, contrasting with monogynous species where queen removal typically extends lifespan. Workers in single-queen colonies lived longer and were more fertile than in two-queen colonies. Queen number did not directly affect oxidative stress resistance or fat body transcription, though the latter depended on an interaction with worker task. Furthermore, younger nurses demonstrated higher fecundity, oxidative stress resistance and upregulated antioxidant genes compared to older foragers. Absent or minor shifts in fecundity and transcription with queen age, respectively, indicated physiological stability with age. Our research highlights distinct caste- and tissue-specific ageing patterns in this supercolonial species, deviating from typical monogynous ants.

## Introduction

A fundamental tenet of life history theory predicts that organisms are confronted with trade-offs. These trade-offs can be driven by resource allocation, e.g., organisms must decide on whether to invest more resources into body maintenance or reproduction. They can also evolve as a consequence of suboptimal gene regulatory networks for both, reproduction and body maintenance with age (Kirkwood, 2017; Maklakov and Chapman, 2019). Ageing – the unavoidable intrinsic deterioration with age, limits the reproductive potential of an organism, often resulting in a negative correlation between longevity and fecundity (Chapuisat and Keller, 2002; Partridge and Harvey, 1988). As organisms age, the accumulation of molecular and cellular damage leads to functional decline, resulting in increased risk of disease and death with advancing age (Kirkwood et al., 1997). The manifestation of reactive oxygen species (ROS) and the organism’s fight to neutralize its damage, results in oxidative stress, a proximate cause of ageing. Metabolically costly activities such as reproduction may favour the production of ROS (Kramer et al., 2021; Selman et al., 2012). Nevertheless, if organisms invest into antioxidants, ROS can be neutralized before critical damage occurs (Münch et al., 2008; Ray et al., 2012; Seehuus et al., 2006).

Social insects, such as ants, termites, some bees and wasps, represent fascinating models to investigate the evolution of senescence. Obligate reproductive division of labour has led to the emergence of different phenotypes with extremely divergent morphologies, behaviours, lifespans, and fecundities. Social insect queens are not only extremely fertile, but also the longest-lived individuals in the colony, whereas workers are typically short-lived and sterile (Keller, 1998; Kramer et al., 2016). Thus, social insects show an apparent reversal of the trade-off between longevity and fecundity. One reason for this may be that social insect queens are naturally supplied with ample resources by their colony. While the molecular mechanisms underlying caste-specific ageing in social insects are not fully resolved, it is known that queens invest more into body repair and maintenance, presumably promoting their extended lifespans (several decades in ants and termites) (Lin et al., 2021; Negroni et al., 2019). Investment and a physiological focus on maintaining the body, growth and reproduction are, therefore, key factors determining ageing and fertility.

The lifespan of queens is also influenced by environmental and social factors. In ants, the average life expectancy of a queen is much higher in species with single-queen societies than in polygynous species with several queens (Keller and Genoud, 1997). The social organisation of the colony and queen founding behaviour are often correlated. Queens of monogynous species are well provisioned and often found their colonies independently, whereas queens of polygynous species typically return to the natal nest. Thereafter, they often establish colonies in collaboration with other queens and workers. Queen loss in monogynous societies ultimately leads to the demise of the colony and impacts worker behaviour and physiology. In contrast, the loss of one queen in a polygynous society has often only minor consequences for the colony as queens can easily be replaced (Keller, 1998).

Under queenless conditions, workers of many ants develop their ovaries, fight over reproductive dominance and lay haploid, male-destined eggs (Boomsma, 2009; Heinze et al., 1997). This option is not available to workers from all ant species as worker sterility is found in invasive species and other species with large colonies, such as *Atta* leaf-cutting ants (Aron et al., 2001; Dijkstra and Boomsma, 2006). As ant queens emit pheromones that indicate their presence and fecundity (Holman, 2018; Oliveira et al., 2020), worker reproduction under queenright conditions is rare (Bourke, 1988; Foitzik and Herbers, 2001). The onset of egg production in queenless workers has a strong impact on their physiology and immunity, as they become more resistant to oxidative stress, exhibit altered transcriptional activity in their brain and fat body, and often live longer than workers in queenright colonies (Kohlmeier et al., 2017; Lopes et al., 2020; Majoe et al., 2021; Negroni et al., 2021). Thus, worker fecundity seems to be a highly plastic trait, which is positively linked to lifespan (Heinze and Schrempf, 2008). The likelihood that ant workers develop their ovaries is also influenced by their age and role within the colony. In social insects, younger workers typically remain in the nest to care for the brood and queen, while older workers take on riskier tasks outside the nest, such as foraging or nest defence (Heinze and Schrempf, 2008; Tofilski, 2002; Wilson, 1980). This age polyethism causes worker age and behaviour to be closely intertwined. Under queenless conditions, younger inside workers are more likely to activate their ovaries and start reproducing compared to older outside workers (Bourke, 1988).

In comparison to species where workers have functional ovaries, our understanding of how species with sterile ant workers respond to queen loss is limited. As sterile workers cannot produce male offspring, investing in reproduction is not an option. For such species, workers possess ovaries and can even lay eggs, but these eggs never develop, and are considered functionally sterile. For example, the loss of the queen in the unicolonial ant *Lasius neglectus* did not affect the susceptibility to oxidative stress and gene expression of workers (Majoe et al., 2024). Using a similarly invasive, polygynous species with regular queen replacement and functionally sterile workers, with developed ovaries might help us to tease apart whether the extended lifespan in species with functional worker ovaries is due to having developed ovaries, or, due to their ability to reproduce.

Our study system, the dolichoderine ant species *Tapinoma magnum* fulfils these requirements and is a great study system to investigate this question. *T. magnum* belongs to the West and Central Mediterranean *Tapinoma nigerrimum* complex and has the widest distribution with the strongest invasive potential (Dekoninck et al., 2016; Seifert et al., 2017). In its invasive range, this ant species is highly polygynous and supercolonial with colonies containing thousands of workers and up to 350 queens. Most mated young queens stay in or near their natal colony and become adopted into neighbouring nests (Seifert et al., 2017). Workers exhibit a size polymorphism and have well-developed ovaries (Seifert et al., 2017). To our knowledge, worker-laid eggs in this species never develop into males and are likely non-viable trophic eggs (Gobin et al., 1998; Gobin and Ito, 2000).

In this study, we investigated how queen number affects worker lifespan, physiology, and gene expression as well as the physiological and molecular influence of age on workers and queens. In contrast to previous studies that examined the influence of queen presence in mainly monogynous or facultatively polygynous species (Majoe et al., 2021; Negroni et al., 2021), we tested how workers respond to queen loss, monogyny, and polygyny. These studies uncovered a positive link between worker survival and fecundity that were associated with higher investment in antioxidants, detoxification enzymes and immunity. Here, we investigated whether queen number similarly leads to such changes in survival and fecundity, as well as to transcriptional changes in the fat body, which plays an essential role in energy storage, growth, and reproduction (Arrese and Soulages, 2010; Cervoni et al., 2017). Moreover, since *T. magnum* workers can lay trophic eggs, we hypothesized that worker age influences fecundity and survival. We expected younger inside workers to possess a higher reproductive potential, linked to an increased oxidative stress resistance compared to the older outside workers.

Similarly, age can lead to changes in fecundity and gene expression of ant queens (Negroni et al., 2019; Von Wyschetzki et al., 2015). Since queen lifespan is often linked to the social organisation of the species (Keller and Genoud, 1997), we expected *T. magnum* queens to be shorter-lived (∼1-2 years) and possibly age differentially than long-lived (> 15 years) queens of monogynous species. It has been shown that fecundity can be maintained throughout a queen’s life and only decreases shortly before death in short-lived *Cardiocondyla obscurior* queens, while old, long-lived *Temnothorax rugatulus* queens switch from investment in immunity to the production of antioxidants (Jaimes-Nino et al., 2022; Negroni et al., 2019). Thus, we focused on age-related changes in gene expression and fecundity between relatively young (≤ 3 months old) and old (≥ 16 months old) *T. magnum* queens.

## Material and Methods

### Study site, collection, and laboratory maintenance

We collected ants from a supercolony of *Tapinoma magnum* on a strip of fallow land, formerly a plant nursery, in Ingelheim am Rhein, Germany (49°58’39.8“N, 8°03’17.3”E) in June and July 2020, and in June 2021. In 2020, 13 dealate queens, a few thousand workers and brood were collected. The collection trips in 2021 occurred during the nuptial flight, resulting in the collection of 13 alate queens, over 30 males, 50 dealate queens, and a few thousand workers and brood.

The colonies were kept in a 25°C climate chamber at the Johannes Gutenberg University Mainz. Colonies from the two collection years were housed in separate boxes (78 cm x 56 cm x 18 cm), which were coated with Fluon® (Whitford GmbH, Diez, Germany) to prevent the ants from escaping. Each box consisted of two to three artificial nest sites built out of two Petri dishes glued together and connected by a 1 cm hole, which was tightly plugged with a piece of cotton. The lower Petri dish (7.5 cm diameter, 4 cm high) was filled with water and the top Petri dish (9.5cm diameter, 1.2cm high) had a loose lid with holes, functioning as a nest entrance. Each nest was covered with a plastic flowerpot (13cm diameter, 12cm high) to create a dark environment for the ants. The colonies were fed twice weekly with a food mix containing honey, eggs, and vitamins and once a week with crickets. Queens collected in 2020 were maintained under similar conditions for at least 16 months. Alate queens collected in 2021 were kept with males to fertilise them. After copulation, the young queens shed their wings and became reproductive.

### Experimental set-up

To investigate the effect of queen number on worker survival, fecundity, and gene expression, we separated the source colony collected in 2021 into 10 sets with each set consisting of three treatment boxes. We randomly assigned workers to one of three treatments: no queen present (“queenless” treatment), one queen present (“monogynous” treatment), and two queens present (“polygynous” treatment). Inside workers were collected near or on the brood pile, while outside workers were collected in the foraging arena outside the nest. Each sub-colony contained 176 inside workers and 176 outside workers with a 1:1 ratio of large and small workers, including a similar amount of brood provided in each sub-colony (Fig. 1). Pupae were removed before emergence. The ants were maintained in their treatments for 58 days and worker mortality was documented twice per week by counting the number of dead workers within each experimental box. After the 58 days, large inside and outside workers were frozen at −80°C for later ovary (fecundity analyses) and fat body dissections (RNAseq analyses).

**Figure 1:**
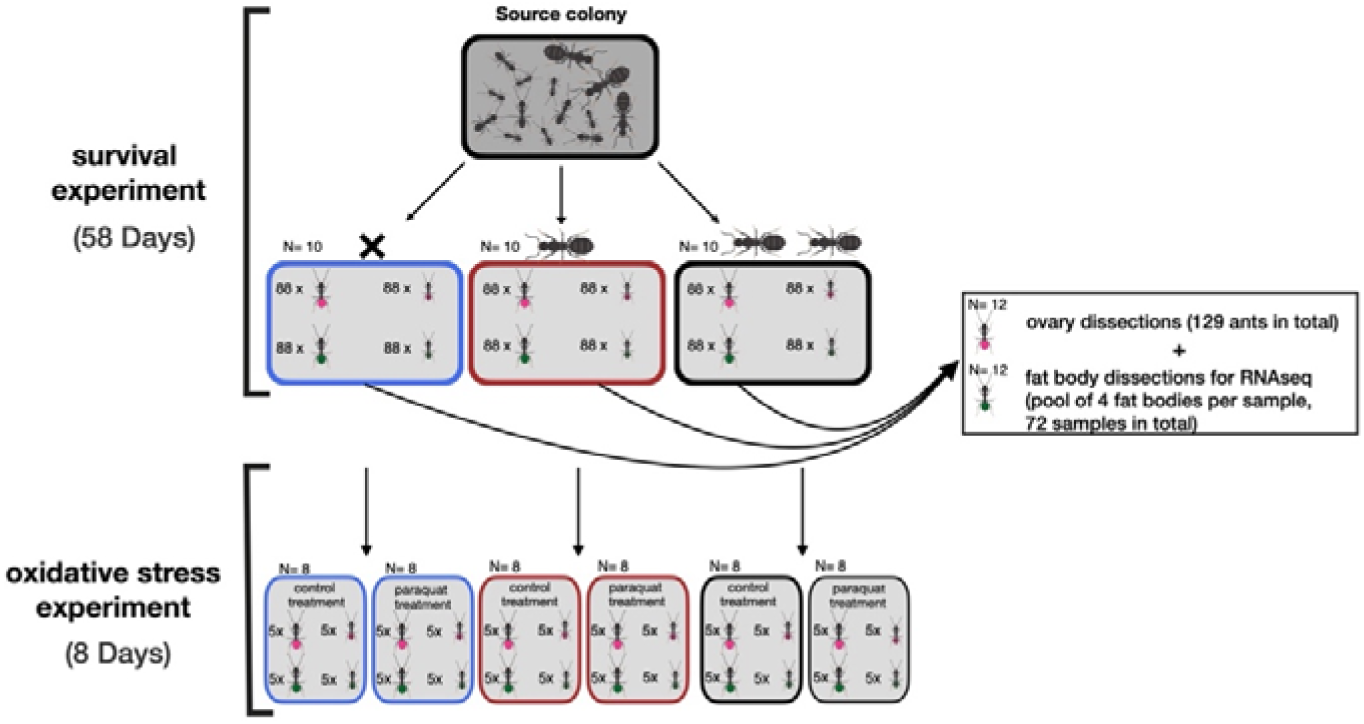
Experimental set-up. For the survival experiment, the source colony was divided into three sub-colonies (N= 10) consisting of no queen (queenless treatment, blue box), one queen (monogynous treatment, red box) and two queens (polygynous treatment, grey box). The sub-colonies consisted of 352 workers comprising of similar numbers of large/small inside and outside workers. Worker mortality was documented twice per week for 58 days. For the fecundity and RNAseq analyses, large inside (pink, N= 12) and outside (green, N= 12) workers from the three treatments were used. The remaining large and small inside and outside workers were subjected to either an oxidative stress treatment or control treatment for eight days.

Queens collected in 2020 were at least 16 months old when sampled, but likely older as they were already without wings at collection. They were afterwards classified as “old” in our transcriptome-based analysis. The winged unmated queens collected in 2021 were not older than three months when sampled and were classified as “young”. After the young queens became reproductive, we created 12 sub-colonies with each consisting of one young and old queen with tending workers (24 queens in total). We marked the queens with thin wires of different colours to denote their age cohort. Each sub-colony consisted of a mix of large and small inside and outside workers from both collection years (200 workers per sub-colony). The queens were maintained in these standardized sub-colonies for 14 days and, thereafter, frozen at −80°C for later ovary, brain, and fat body dissection for fecundity and RNAseq analyses.

### Oxidative stress experiment

After the 58-day survival experiment, we took 10 large and 10 small inside workers as well as 10 large and 10 small outside workers from each treatment and colour-marked the workers with paint markers (Edding® 750; 960 ants in total, 8 sets). We randomly assigned five large and five small inside workers, five large and five small outside workers to either the control or the oxidative stress treatment (Fig. 1). The next day, workers assigned to the oxidative stress treatment (480 total number of workers per treatment) were subjected to a 0.46 M solution of Paraquat dichloride (CH_3_(C_5_H_4_N)_2_CH_3_・2Cl) dissolved in Millipore water. Paraquat is an herbicide inducing oxidative stress through superoxide formation (Cousin et al., 2013). We applied the paraquat solution twice to each worker’s head with a size 1 Vernissage^TM^ paintbrush. We covered the entire top of the head so that the amount was approximately proportional to that ants’ body size. After application, each ant was isolated for two hours in a 1.5 ml Eppendorf tube to avoid the transfer of the solution to other ants via trophallaxis (mouth-to-mouth fluid exchange), and to facilitate self-grooming and ingestion. Workers of the control treatment were treated similarly but here we applied pure Millipore water instead of the paraquat solution. These treatments were repeated every two days over a period of eight days and worker survival was monitored three times per day for eight days (Majoe et al., 2021).

### Fecundity analyses

To investigate whether worker ovarian development depended on their location (used as a proxy for age) and queen number, we dissected the ovaries of large inside and outside workers (Fig. 1). Ovaries were dissected under a stereomicroscope (Leica S9i, Microsystems CMS GmbH, Wetzlar, Germany) and pictures were taken using Leica LAS software (version 4.12.0). For each ant, ovariole length was measured with ImageJ2 Fiji software (version 2.9.0) and the mean of the two longest ovarioles was calculated. The number of developing oocytes was counted as well. All queen ovaries were dissected and the mean ovariole length was measured as described for the workers. The presence of a sperm-filled spermatheca could be confirmed for all queens.

### Statistical analyses

The coxme package (version 2.2-18.1; Therneau, 2020) in RStudio (R version 4.2.3; R Core Team, 2018) was used to build cox-regression mixed-effect models for both survival experiments (58-day survival and oxidative stress experiment). The variable “queen number” (queenless/ monogynous/ polygynous) was included as fixed factor, and “experimental box” as random factor in the analysis of the 58-day survival experiment. For the oxidative stress experiment, we additionally included the “treatment” (oxidative stress treatment/ control), “worker location” (inside/ outside) and “worker size” (large/ small), as well as their interaction as fixed factors, and “experimental box” as a random factor. Survival within each treatment (queen treatment/ oxidative stress/ control) was assessed independently. Hypothesis testing was performed using the ‘Anova’ function from the package car (version 3.1-2; Fox and Weisberg, 2019). The package ggplot2 (version 3.4.2; Wickham, 2016) was used to plot the Kaplan-Meier survival curves. To compare whether worker survival depended on queen number, pairwise comparisons were performed for: queenless vs. monogynous; queenless vs. polygynous; and monogynous vs. polygynous.

We modelled the mean ovariole length using a linear mixed-effect model from the package lme4 (version 1.1-33; Bates et al., 2015) and tested the effect of the explanatory variables with the ‘Anova’ function (car package). The number of oocytes in development was modelled using a generalized linear mixed-effect model with the package glmm (version 1.4.4; Knudson, 2022) with a ‘Poisson’ distribution and tested with the ‘Anova’ function. For both analyses, “queen number”, “worker location” and their interaction were set as fixed variables with “colony ID” and “experimental box” included as random factors.

The ovarian profile of 11 young and 11 old queens (22 total number of queens) was analysed independently from the workers but similarly using a linear mixed-effect model with the package lme4. “Mean ovariole length” was included as the response variable, “queen age” (young/ old) as a fixed factor and “colony ID” as a random factor. As above, hypothesis testing was performed with the ‘Anova’ function.

### Differential expression analysis

For the differential expression analysis, we used 72 worker samples (fat body) and 24 queen samples (brain and fat body). Detailed information on tissue dissection, RNA extraction, and sequencing can be found in the supplementary material. Raw RNAseq read quality was assessed, and filtering was performed to remove adaptors using Fastp (version 0.2; Chen et al., 2018). Post-filtered sequence quality was assessed using FastQC (version 0.11.8; Andrews, 2010) with the data being of high quality to perform differential expression analyses. Since there is no reference genome assembly available for *Tapinoma magnum,* we generated a *de novo* transcriptome assembly using Trinity (version 2.13.2; Grabherr et al., 2011) with the SuperTranscripts option (Davidson et al., 2017). To generate the transcriptome assembly, we used a combination of the filtered queen and worker reads. To quantify transcript abundance, RSEM (version 1.3.3; Li and Dewey, 2011) was used, resulting in an overall alignment rate of 79.4% ± 1.5 (mean ± sd) for the worker dataset and 76.7% ± 2.5 (mean ± sd) for the queen dataset. One queen brain sample was removed from further analyses due to a low mapping rate of 47%. Differential gene expression analysis was conducted in RStudio (R version 4.2.3). For each dataset, transcript abundances were introduced into estimated count matrices with the package tximport (version 1.26.1; Soneson et al., 2015), serving as input for differential gene expression analyses using the DESeq2 package (version 1.38.3; Love et al., 2014). Subsequent transcriptome-based analyses were conducted separately for the worker and queen dataset.

For the worker dataset, contigs with zero reads in at least 11 of the 72 samples were removed from the count matrix, resulting in 73,277 contigs retained in the differential expression analysis. To identify genes whose expression was affected by the interaction between the number of queens and the collection location, we first compared a full model consisting of “colony ID”, “queen number”, “worker location” and the interaction between “queen number” and “worker location” to a reduced model that included “colony ID”, “queen number” and “worker location”. To test the main effect of queen number, we compared a full model containing “colony ID”, “queen number”, and “worker location” to a reduced model consisting of “colony ID“ and “worker location” only. Second, to examine the main effect of worker location on gene expression, we compared a full model containing “colony ID”, “queen number”, and “worker location” to a reduced model consisting of “colony ID “and “queen number” only. To identify differentially expressed genes with similar expression patterns, we conducted a cluster-based analysis with the R package DEGreport (version 1.34.0; Pantano, 2017).

For the queen-based analysis, we ran separate analyses for the two tissues (brain and fat body). Only contigs with more than zero reads in at least five of six samples per tissue were kept, resulting in 55,349 contigs being used for the differential expression analysis. Our full model included “age” and the reduced model consisted of the intercept. All model comparisons for the queen and the worker data were conducted using Likelihood Ratio Tests (LRT) as implemented in DESeq2 with a Benjamini-Hochberg adjusted p-value of 0.05 used as a threshold to determine whether a gene was significantly differentially expressed.

### Gene annotation and functional enrichment analyses

To annotate the Trinity-assembled transcriptome, we first conducted a BlastX homology search against the non-redundant invertebrate protein database (Altschul et al., 1990; downloaded July 2023). Nucleotide sequences were translated into predicted amino-acid sequences using TransDecoder (version 5.7.0-Perl-5.30.0; Haas, 2017), which were then used as input for InterProScan (version 5.54-87.0; Blum et al., 2021) for the identification of conserved functional domains and assignment of associated Gene Ontology (GO) terms to our *de novo* transcriptome assembly. We additionally used OrthoFinder (version 2.5.4; Emms and Kelly, 2019) to examine potential homologues between the closely related Dolichoderine ant *Linepithema humile* (Ensembl Metazoa Biomart) and *Tapinoma magnum*, resulting in the identification of 5,859 potential homologues, of which 63% could be assigned additional GO terms based on homology to *L. humile*. We combined both sets of GO terms to create a GO term database, which was then used in our GO term enrichment analysis. To identify which terms attributed molecular functions functionally enriched within workers and queens, we performed individual GO term enrichment analyses based on the differentially expressed contigs using Fisher’s exact tests (threshold of significance: p < 0.05) implemented by the package TopGo (version 2.50.0; Alexa and Rahnenführer, 2010) using the weight01 algorithm. The scripts used for these analyses were modified versions of those developed by Colgan et al. (2019).

## Results

### Queen number influenced worker survival, while worker location was associated with worker fecundity

To investigate the effect of queen number on worker survival, we monitored worker survival in queenless, monogynous, and polygynous colonies over 58 days. We found that queen number influenced worker survival, as workers from monogynous sub-colonies survived best, followed by workers of polygynous colonies, and then workers of queenless colonies (queen number 0 vs. 1: X^2^= 155, p< 0.001; 0 vs. 2: X^2^= 41.6, p< 0.001; 1 vs. 2: X^2^= −51.7, p< 0.001; Fig. 2A). We further examined the influence of queen number and worker location on worker fecundity by measuring ovariole length and counting the number of developing oocytes in inside and outside workers from each queen treatment. The workers had a mean ovariole length of 1.87 mm (± 0.4 mm sd), which was independent of queen number and worker location (LMER_queen number_: X^2^ = 2.04, p= 0.36; LMER_worker location_: X^2^ = 1.08, p= 0.29; Supplementary Figure S1). In contrast, inside workers had more developing oocytes than outside workers (GLMER: X^2^= 18.76, p< 0.0001; Fig. 2B). Moreover, workers of monogynous sub-colonies exhibited a higher number of oocytes than workers of polygynous sub-colonies (GLMER: X^2^ = 4.88, p= 0.027; Fig. 2B).

**Figure 2:**
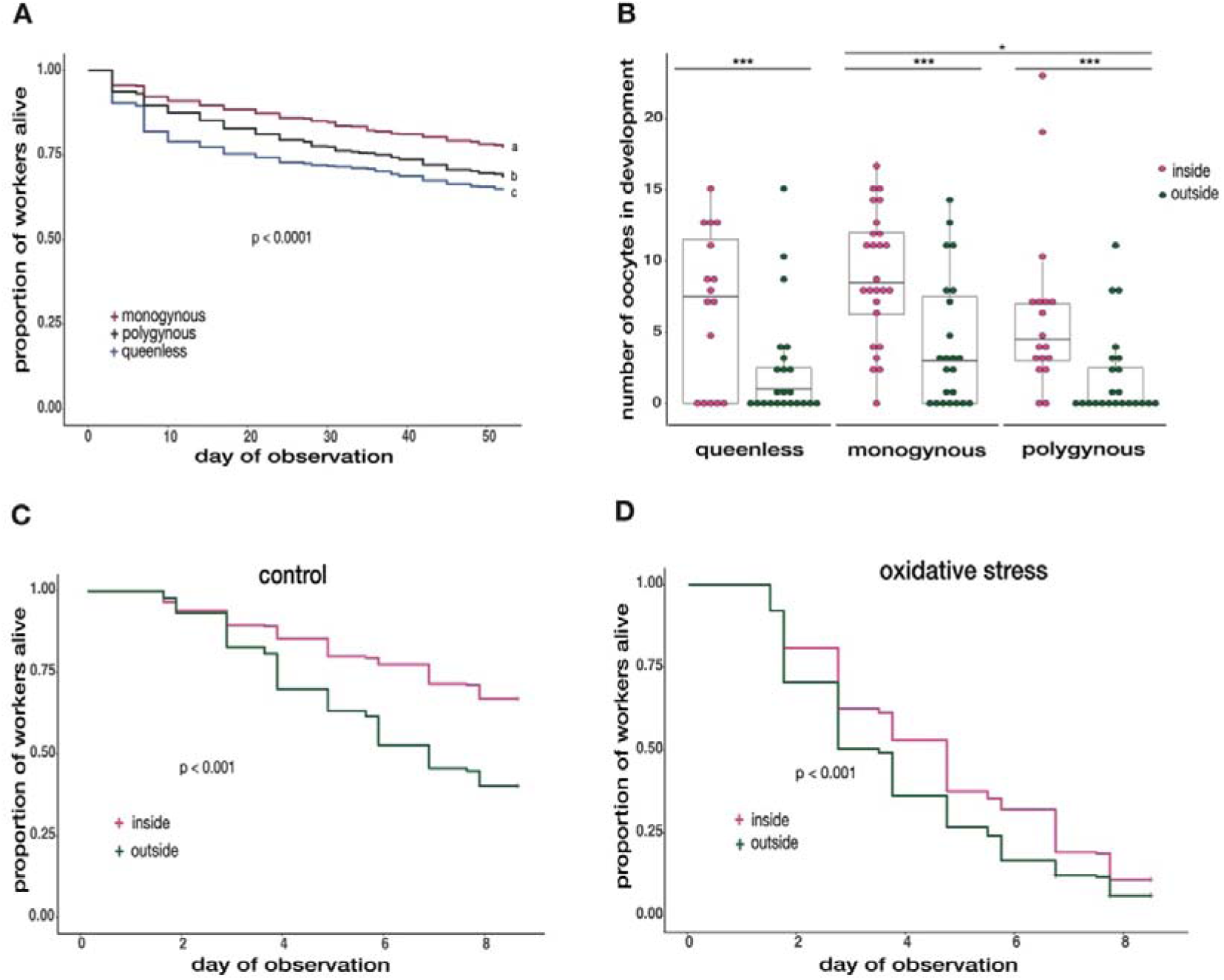
Worker survival, fecundity, and oxidative stress resistance dependent to queen number and worker location. A) Worker survival over 58 days under queenless, monogynous and polygynous conditions. Workers of monogynous (red line) colonies survived best, followed by workers of polygynous (grey line) colonies and lastly workers of queenless (blue line) colonies (Queen number 0 vs. 1: X^2^= 155, p< 0.001; 0 vs. 2: X^2^= 41.6, p< 0.001; 1 vs. 2: X^2^= −51.7, p < 0.001). B) The number of oocytes dependent to queen number and worker location. Inside (pink) workers had more oocytes in development than outside (green) workers (GLMER: X^2^= 18.76, p< 0.0001). Workers of monogynous colonies developed more oocytes than workers of polygynous colonies (GLMER: X^2^= 4.88, p= 0.027). C) Worker survival within the control treatment from the oxidative stress experiment. Inside (pink line) workers have a higher survival probability than outside (green line) workers (X^2^= 37.78, p< 0.0001). D) Worker survival under oxidative stress treatment. Inside workers show a higher resistance to oxidative stress than outside workers (X^2^= 26.81, p< 0.0001).

### Worker location but not queen number influenced worker resistance to oxidative stress

To test whether queen number, worker location, and worker size influenced the resistance of workers to oxidative stress, we subjected large and small inside and outside workers from each queen treatment to either a paraquat-induced oxidative stress treatment or the control treatment (Millipore water). The oxidative stress treatment reduced worker survival (X^2^= 307.72, p< 0.0001), while queen number did not influence worker survival under oxidative stress (X^2^= 2.49, p= 0.287). In the control treatment, we found weak evidence that queen number might influence worker survival (X^2^= 4.79, p= 0.091). Closer examination revealed that this trend followed a similar pattern as to what we observed after our 58-day survival experiment (queen number 0 vs. 1: X^2^= 3.09, p= 0.08; 0 vs. 2: X^2^= 0.002, p= 0.97; 1 vs. 2: X^2^= 3.57, p= 0.058). Inside workers survived better than outside workers in both, the control, and the oxidative stress treatment (control: X^2^= 37.78, p< 0.0001; oxidative stress: X^2^= 26.81, p< 0.001, Fig. 2C/D). Additionally, large workers survived better under oxidative stress than small workers (X^2^= 46.11, p< 0.001), while we found very weak evidence that worker size influenced survival within the control treatment (X^2^= 2.69, p= 0.10; Supplementary Figure S2).

### Worker location rather than queen number influenced fat body gene expression in workers

To investigate the effect of queen number and worker location on fat body gene expression, we analysed expression differences between inside and outside workers from each treatment (queenless/ monogynous/ polygynous). We found no evidence that queen number affected fat body gene expression (0 differentially expressed genes, all BH-adjusted p-value > 1), while worker location was linked to strong transcriptomic differences. Between inside and outside workers, we identified 4,305 significantly differentially expressed genes (DEGs; BH-adjusted p< 0.05). Of these, 2,188 showed an elevated expression in inside workers, while 2,117 DEGs were elevated in outside workers. We identified 108 DEGs, whose expression was affected by an interaction between worker location (inside/ outside) and queen number (queenless/ monogynous/ polygynous), of which, 102 were grouped into three co-expression clusters (Fig. 3A). The first cluster consisted of 31 genes whose expression in inside workers decreased with increasing number of queens. Within outside workers the expression was lower in queenless colonies and similarly increased in monogynous and polygynous colonies. Cluster 2 consisted of 36 genes where expression was highest in inside workers sampled from queenless colonies, while, similar to cluster 1, the expression was decreasing with queen number. Within outside workers, the difference in expression between queenless and monogynous colonies was even stronger compared to cluster 1. The expression in monogynous and polygynous colonies was similarly high. The blast homology search identified two genes in cluster 1 (*serine protease inhibitor ¾-like isoform X3,* LRT: BH-adjusted p= 0.048; *group XIIA secretory phospholipase A2*, LRT: BH-adjusted p= 0.049) and two genes in cluster 2 (*ribosome biogenesis protein WDR12 homolog,* LRT: BH-adjusted p= 0.049; *cell division cycle protein 123 homolog isoform X1,* LRT: BH-adjusted p= 0.037). Finally, cluster 3 contained 35 genes with expression patterns opposite to the first two clusters (Fig. 3A). Within inside workers, expression profiles increased with queen number, while outside workers showed the highest expression in queenless colonies, which was decreasing in monogynous and polygynous colonies. Here, we found nine genes related to immunity and detoxification (e.g., *xanthine dehydrogenase 1-like*, LRT: BH-adjusted p= 0.037; *glucose dehydrogenase*, LRT: BH-adjusted p= 0.047; Kim et al., 2001; Lee et al., 2005), three genes involved in the Ras signalling pathway (e.g., *ras-like GTP-binding protein Rho1 isoform X2*, LRT: BH-adjusted p= 0.013), and *juvenile hormone epoxide hydrolase 1-like* (LRT: BH-adjusted p= 0.047).

**Figure 3:**
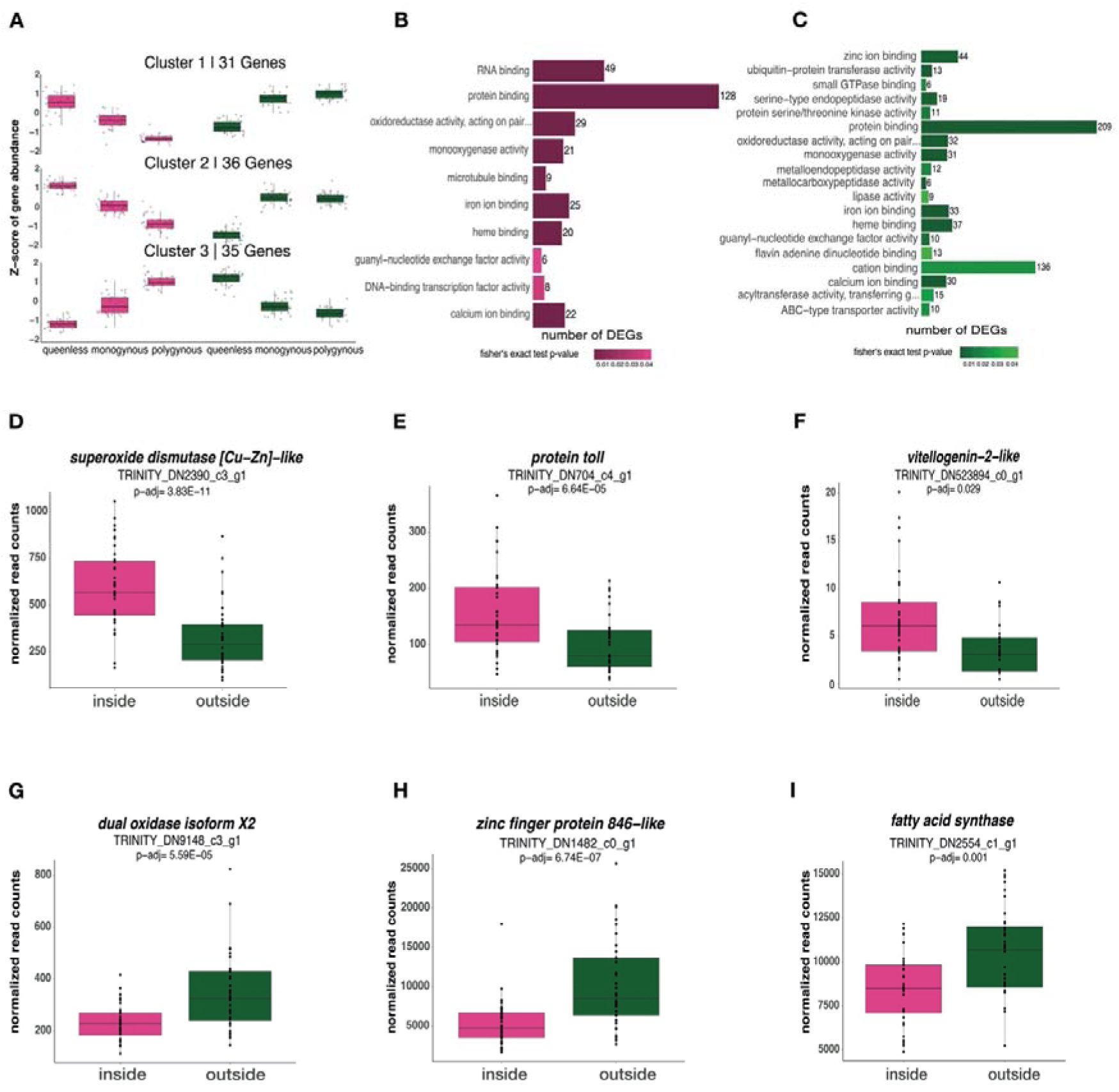
The influence of queen number and worker location on fat body gene expression. A) 102 out of the 108 DEGs from the interaction term (queen number : worker location) were grouped into three clusters based on the expression profiles within each worker caste (pink= inside workers; green= outside workers) and social environment. B) The 10 most enriched GO terms (molecular function) in the list of genes upregulated in inside workers and C) the 19 most enriched GO terms (molecular function) in the list of genes upregulated in outside workers. The size of each bar correlates with the number of DEGs associated with each term, while the colour correlates with the p-value (the darker the colour, the lower the p-value). D)-I) Selected DEGs with notable function within inside workers and outside workers.

Genes with higher expression in inside workers were enriched for 10 Gene Ontology (GO) terms (Fig. 3B, Supplementary Table S1). Among these, the most significantly enriched terms were *oxidoreductase activity* (Fisher’s exact p< 0.0001)*, iron ion binding* (Fisher’s exact p< 0.0001), and *monooxygenase activity* (Fisher’s exact p< 0.0001). In comparison, we identified 19 enriched GO terms for the DEGs upregulated in outside workers. Next to terms also represented in inside workers, we found some terms related to lipid metabolic processes (e.g., lipase activity, Fisher’s exact p= 0.036) and proteolysis (e.g., metallocarboxypeptidase activity, Fisher’s exact p< 0.001), drawing a more diverse expression profile of in outside workers (Fig. 3C; Supplementary Table S2).

Within the DEGs upregulated in inside workers, our blast homology search identified the antioxidants *superoxide dismutase [Cu-Zn]-like* (LRT: BH-adjusted p= 3.83E-11, Fig. 4D), *transferrin* (LRT: BH-adjusted p= 0.003), as well as *protein toll* (LRT: BH-adjusted p= 6.64E-05, Fig. 4E). Moreover, we found *vitellogenin-1* (LRT: BH-adjusted p= 0.045)*, vitellogenin-1-like* (LRT: BH-adjusted p= 0.012), and *vitellogenin-2-like* (LRT: BH-adjusted p= 0.029, Fig. 4F; all *conventional vitellogenins*, *C-Vg;* Kohlmeier et al., 2018). Within the genes upregulated in outside workers, we found higher expression of *dual oxidase isoform X2* (LRT: BH-adjusted p= 0.035, Fig. 4G), 22 *zinc finger proteins* (e.g., *zinc finger protein 846-like*, LRT: BH-adjusted p= 6.74E-07, Fig. 4H), 24 DEGs involved in lipid metabolism (e.g., *fatty acid synthase*, LRT: BH-adjusted p= 0.001, Fig. 4I), as well as four odorant receptors (e.g., *odorant receptor coreceptor*, LRT: BH-adjusted p= 1.08E-11).

**Figure 4:**
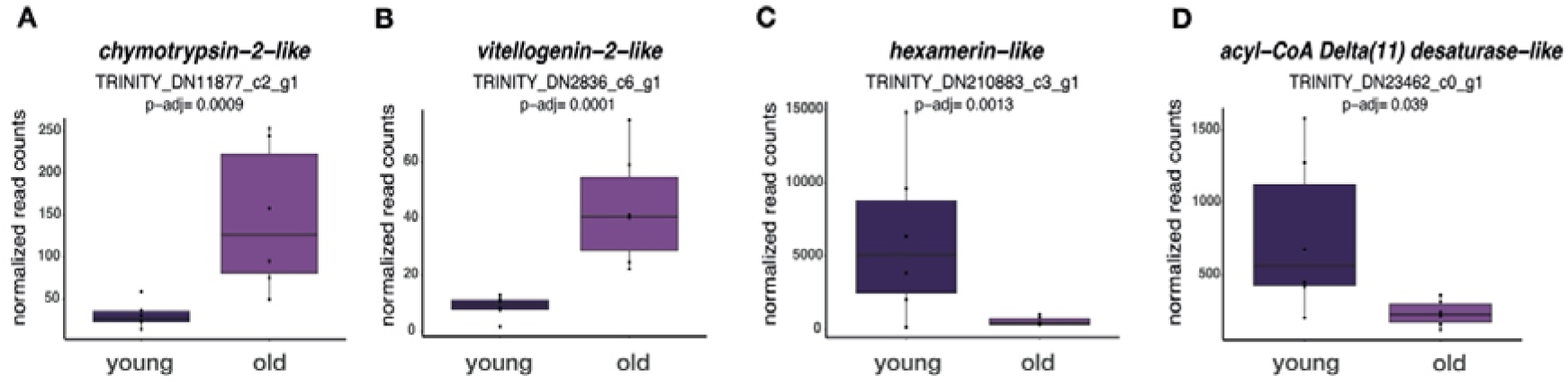
Candidate DEGs of interest differing in expression in the fat body between young and old queens. **A)** Normalized read counts of *chymotrypsin-2-like* and **B)** *vitellogenin-2-like* which are significantly overexpressed in old queens. **C)** Normalized read counts of *hexamerin-like* and **D)** *acyl-Coa Delta(11) desaturase-like* which are overexpressed in young queens.

### Queen age elicited no changes in fecundity but minor transcriptional shifts in fat body gene expression

To investigate the effect of age on queen fecundity and gene expression, we measured ovariole length and analysed brain, as well as fat body gene expression of young and old *T. magnum* queens. Age did not influence ovarian development as young queens had similarly long ovarioles as old queens (LMER: X^2^= 0.02, p= 0.811; Supplementary Figure S3). Furthermore, we did not detect any age-related transcriptomic changes in the brain, while we identified 126 DEGs in the fat bodies between young and old queens. Of these, 75 DEGs were upregulated in old queens, while 51 DEGs were upregulated in young queens. In old queens, we found an overexpression of *chymotrypsin-2-like* (LRT: BH-adjusted p= 0.0009, Fig. 4A), *vitellogenin-1-like* (LRT: BH-adjusted p= 0.005), and *vitellogenin-2-like* (Fig. 4B; LRT: BH-adjusted p= 0.0001, *C-Vg;* Kohlmeier et al., 2018). Young queens showed increased expression of *hexamerin-like* (LRT: BH-adjusted p= 0.0013, Fig. 4C) and *acyl-CoA Delta(11) desaturase-like* (LRT: BH-adjusted p= 0.039, Fig. 4D). We could not identify significantly enriched GO terms within the DEGs between young and old queens.

## Discussion

### Physiological and transcriptional effects of queen number and worker location

We investigated whether queen number and worker location affected worker survival, fecundity, oxidative stress resistance, and fat body gene expression in the invasive ant *Tapinoma magnum*. Our two-month survival experiment revealed that workers of monogynous colonies survived best, followed by workers of polygynous colonies, with the lowest survival in workers of queenless colonies. Workers of monogynous colonies had more oocytes in development compared to those from polygynous colonies, while inside workers had more oocytes than outside workers. Worker resistance to oxidative stress and gene expression of the fat body were not influenced by queen number whereas worker location strongly influenced both.

Our results contrast with previous studies on other ant species (Kohlmeier et al., 2017; Majoe et al., 2021), as *T. magnum* workers showed the highest mortality in queenless colonies. Queens of insect societies are normally the only group members that reproduce. The presence of a queen regulates essential aspects of worker behaviour and physiology, either via direct behavioural interactions or indirectly via pheromones (Breed and Gamboa, 1977; Holman et al., 2010). Queen loss affects, to some extent, the whole colony (Brunner and Heinze, 2009; Stroeymeyt et al., 2007). To our knowledge, there is no evidence for increased worker reproduction after queen loss in strictly polygynous ant species (Friend and Bourke, 2014). In fact, our results indicate that queen loss decreases worker survival, without influencing worker fecundity. In polygynous species, losing all queens is extremely unlikely and may have been too stressful for the workers, possibly increasing worker mortality.

The higher survival in monogynous colonies might be linked to the increased oocyte development and presumably increased trophic egg-laying in workers kept in monogynous compared to polygynous conditions. Trophic eggs provide essential nutrients, especially proteins, for growth and reproduction in larvae and queens (Gobin et al., 1998; Gobin and Ito, 2000). In monogynous colonies, egg production hinges on the sole queen, while queen fecundity and nutritional care provided to queens are correlated (Hannonen et al., 2002; Trettin et al., 2011; Chen and Vinson, 2000). In the Argentine ant *L. humile*, queens may receive less food under polygynous conditions compared to monogynous conditions (Keller, 1988). Thus, the increased production of oocytes in monogynous workers followed by increased trophic egg-laying might be linked to a higher nutritional care provided to the single queen. Workers might have increased their own lifespan to increase the fitness and survival of the single queen.

In contrast, queen number had no influence on worker survival under oxidative stress, while we found weak evidence that workers survived better in the control when they had previously been held in monogynous conditions. Therefore, the strong effect of the paraquat-induced oxidative stress treatment may have covered any potential effects of queen number. Main differences in oxidative stress resistance were driven by worker location, supporting our hypothesis that inside workers were younger and had a higher investment in body maintenance. For older outside workers, such investment might not be worthwhile for the relatively shorter remaining lifespan (Kohlmeier et al., 2017; Majoe et al., 2021). Moreover, our transcriptomic analysis revealed that inside and outside workers responded with divergent transcriptional shifts to queen number. Genes of interest in the first cluster were *serine protease inhibitor ¾-like isoform X3* and *group XIIA secretory phospholipase A2*. Serine protease inhibitors are involved in immune responses (Kanost, 1999; Kanost and Clarke, 2005), while the inhibition of phospholipases A2 (PLA_2_) impairs egg-laying behaviour, metabolism, and immunity in insects (Stanley and Kim, 2019). Inside workers upregulated these genes under queenless conditions the most, while outside workers expressed these genes the least in queenless colonies. In cluster 2, genes of interest were related to detoxification (*ribosome biogenesis protein WDR12 homolog*) and the cell division cycle (*cell division cycle protein 123 homolog isoform X1*). Cell cycle division proteins are involved in various biological processes and are often increased after injuries (Pal and Raj, 2022). Thus, young inside workers might have responded more strongly to the queenless environment by an internal battle between the expression of stress-inducing but also immunity-related genes, while older outside workers were likely older and would potentially die earlier independent of queen number. In cluster 3, we found more genes related to immunity and detoxification (e.g., *xanthine dehydrogenase 1-like)*, as well as Ras proteins (e.g., *ras-like GTP-binding protein Rho1 isoform X2*), and *Juvenile hormone epoxide hydrolase 1-like (JHEH)*, upregulated in inside workers within monogynous and polygynous colonies compared to queenless colonies, while outside workers expressed these genes most under queenless conditions. Ras proteins play a central role in cell cycle regulation, wound healing, and tissue repair (Boonstra et al., 1995; Stacey, 2003). *JHEH* plays an important role in the degradation pathways of juvenile hormone (JH). In adult females of *Drosophila melanogaster,* JH is required for oogenesis and reproductive maturation (Dubrovsky et al., 2002). Moreover, gene expression varied less in outside workers within monogynous and polygynous colonies in all three clusters.

Worker location was associated with strong changes in gene expression, likely explaining why inside workers showed a higher resistance to oxidative stress. Young inside workers invested more into antioxidant genes (*superoxide dismutase [Cu-Zn]-like* and *transferrin*), accompanied with an overexpression of *vitellogenin-2-like* (*C-*Vg; Kohlmeier et al., 2018) and *protein toll-like*. Superoxide dismutase and transferrin are known to eliminate reactive oxygen species (ROS) while they are also linked to the individual’s reproductive potential (Nojima et al., 2015; Tasaki et al., 2018). Vg proteins are synthetized in the fat body and secreted into the haemolymph before they are transported into the oocytes (Hagedorn and Kunkel, 1979). Elevated expression of Vg is usually more abundant in queens but can also be found in younger inside workers compared to older outside workers (Corona et al., 2007; Wu et al., 2021), while its expression also correlates with the protection against oxidative stress (Seehuus et al., 2006). Moreover, the insect Toll pathway and its receptors play an important role in both, immunity and development (Evans et al., 2006). Thus, the higher resistance to oxidative stress in combination with the overexpression of antioxidants and increased fecundity in young inside workers implies that in *T. magnum* too, worker age and behaviour are closely intertwined.

In older outside workers, we found the overexpression of *dual oxidase isoform X2*, *zinc finger proteins* and *fatty acid synthase*. Dual oxidase induces oxidative stress to challenge gut microbiome bacteria (Ha et al., 2005; Sistermans et al., 2023), while zinc is involved in regulatory pathways (Kandel, 2009; Klug, 2005). Fatty acid synthase takes part in maintaining energy metabolism, promoting the survival of older animals (Chaudhari and Kipreos, 2018). It might help to compensate the effects of cellular damage of ageing in old outside workers. Moreover, we found four odorant receptors overexpressed in the fat body of outside workers. The role of these proteins within the fat body of insects remains unclear (Chen et al., 2019).

### The physiological and molecular influence of ageing in queens

We investigated the effects of ageing on fecundity and tissue-specific gene expression in young and old *T. magnum* queens. We found no evidence that age influenced fecundity or brain gene expression, while fat body gene expression varied depending to queen age (126 DEGs).

We found a higher expression of *chymotrypsin-2-like, vitellogenin-1-like,* and *vitellogenin-2-like* (C-Vgs; Kohlmeier et al., 2018) in old queens. Chymotrypsin is often expressed as immune response against (bacterial) infection (Viljakainen et al., 2018), possibly linked to an increase in body repair and maintenance within old queens. In *P. barbatus, vitellogenin 1* expression is higher in queens and nurses, while *vitellogenin 2* is higher expressed in foragers (Corona et al., 2013). Thus, these copies might stand in contrast to each other, one indicating a high reproductive potential and the other indicating reproductive senescence within old queens of *T. magnum*.

Young queens overexpressed *hexamerin-like* and *acyl-CoA Delta(11) desaturase-like*. In *Camponotus festinatus* founding queens, high levels of *hexamerin* (larval serum proteins, e.g., storage proteins; Beintema et al., 1994) are related to the production of the first batch of eggs (Martinez and Wheeler, 1994). Supercolonies of *T. magnum* may contain over 350 queens (Seifert et al., 2017). Thus, the workers might not provide each queen with the same amount of food and care (Hannonen et al., 2002; Keller, 1988; Chen and Vinson, 2000), resulting in young queens possibly using up their storage proteins to start reproduction. Acyl-CoA desaturases are involved in various biological processes (Hazel and Eugene Williams, 1990; Miyazaki and Ntambi, 2003) including the production of cuticular hydrocarbons (CHCs). They occur in high levels within *L. humile* and *Solenopsis invicta* and improve the species’ ability to quickly adapt to new ecological niches, possibly favouring their invasive success (Helmkampf et al., 2015).

Our results contrast with a previous study on *Temnothorax* ants showing that older queens have better developed ovaries (Negroni et al., 2019). Indeed, differences in queen ovarian development might depend on the age gap between young and old queens (Seistrup et al., 2023). In our study, old queens were at least one year older than young queens. Queen lifespan varies among species and correlates with the number of queens within the nest (Keller, 1998; Keller and Genoud, 1997). For instance, related species as *Tapinoma sessile* and *L. humile* exhibit queen lifespans within the range of a few weeks to one year (Keller, 1998). Thus, *T. magnum* queens likely exhibit an overall shorter lifespan, potentially explaining the lack of differences we found in their ovary development. Similarly, the lack of age-related changes in fecundity may be a trait of invasive species (Majoe et al., 2024). Compared to previous studies (Negroni et al., 2019; Von Wyschetzki et al., 2015), we did not find many age-related differences in fat body gene expression (∼1500 DEGs vs. 126 DEGs). However, in the unicolonial ant *L. neglectus* similarly low age-dependent shifts in gene expression were found (Majoe et al., 2024; 165 DEGs). The lack of strong age-dependent changes may correlate with the polygynous nature and the corresponding shorter life expectancy of queens in this species, indicating that fecundity and gene expression are maintained throughout their lives (Jaimes-Nino et al., 2022). In fact, we have no exact information on the life expectancy of *T. magnum* queens. Thus, queens might deteriorate faster towards the end of their lives, but our queens might have been middle-aged which is why we could not detect strong age-related changes in gene expression and fecundity (Jaimes-Nino et al., 2022; Majoe et al., 2024).

## Conclusion

Our study provides novel insights into the physiological and molecular responses of the supercolonial ant *Tapinoma magnum* towards different social environments, indicating a clear distinction of life-history traits compared to non-invasive species (Kohlmeier et al., 2017; Majoe et al., 2021; Negroni et al., 2021). We did not only test differences in worker survival between queenless and queenright colonies but also show that workers survived better in monogynous colonies than in polygynous colonies, while removing all queens increased mortality. Further, we show that physiological and transcriptional changes with worker age are comparable to other ant species (Majoe et al., 2021; Negroni et al., 2021), while queen age was not associated with strong changes in fecundity or gene expression. Our findings indicate that *T. magnum* queens are likely overall shorter-lived and maintain fecundity and gene expression throughout their lives (Jaimes-Nino et al., 2022; Keller and Genoud, 1997).

## Author contributions

SF, RL, MM and AL designed the experiment. Ants have been collected by SF, MM, SS, and AL. SS, supervised by SF and co-supervised by AL, conducted the survival and the oxidative stress experiment. AL analysed the data collected by SS, dissected and measured ovaries from workers as well as dissected fat bodies and extracted the RNA. MM and AL dissected brain and fat body from the queens and MM helped AL with the RNA extractions. SF, RL, TJC and MM helped AL in the statistical analyses and interpretation. TJC and AL annotated the transcriptome assembly. AL wrote a first draft and all authors commented on it. AL, SF, RL, TJC finalized the manuscript based on these comments. SF supervised the experimental parts of the project.

## Acknowledgements

We thank Ina Knuf for helping us collect the ants and assisting SS during the experimental phase of the study. We thank Juliane Hartke and Timo Wentong Lin for their help in generating the *de novo* transcriptome assembly. Thanks to Barbara Feldmeyer for her input on the annotation of the *de novo* transcriptome assembly. Moreover, we are grateful to Jenny Fuchs for providing the ant illustrations.

## Data accessibility statement

The raw data and code for the statistical analyses behind the figures are provided in the electronic supplementary material.

Raw read sequences are accessible through the NCBI BioProject ID PRJNA1120839.

## Funding

Funding came from the German Research Foundation via DFG grants FO298/26-1 to SF, and LI3051/3-1 to RL. The study was further supported by GRK2526/1 – Project No. 407023052.

## Notes

### Competing Interest Statement

The authors have declared no competing interest.

