## Supplementary Figure S1 for "Queen number shapes worker longevity, fecundity and gene expression in the invasive, highly polygynous ant *Tapinoma magnum*"

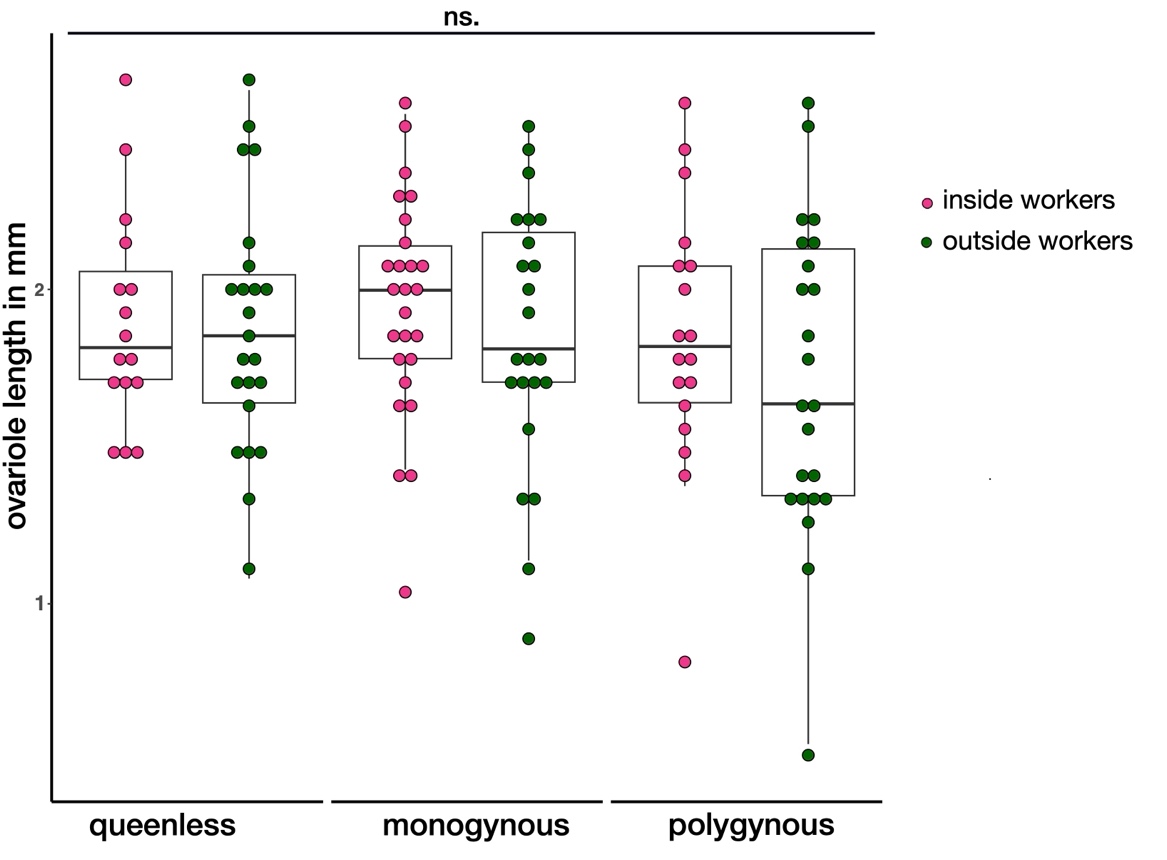

Figure S 1: Mean ovariole length within inside and outside workers of the three queen treatments (queenless, monogynous, polygynous). Inside and outside workers had similarly long ovarioles, independent to queen number (LMER_queen number_: X^2^ = 2.04, p = 0.36; LMER_worker location_: X^2^ = 1.08, p = 0.29).
