## Supplementary Figure S2 for "Queen number shapes worker longevity, fecundity and gene expression in the invasive, highly polygynous ant *Tapinoma magnum*"

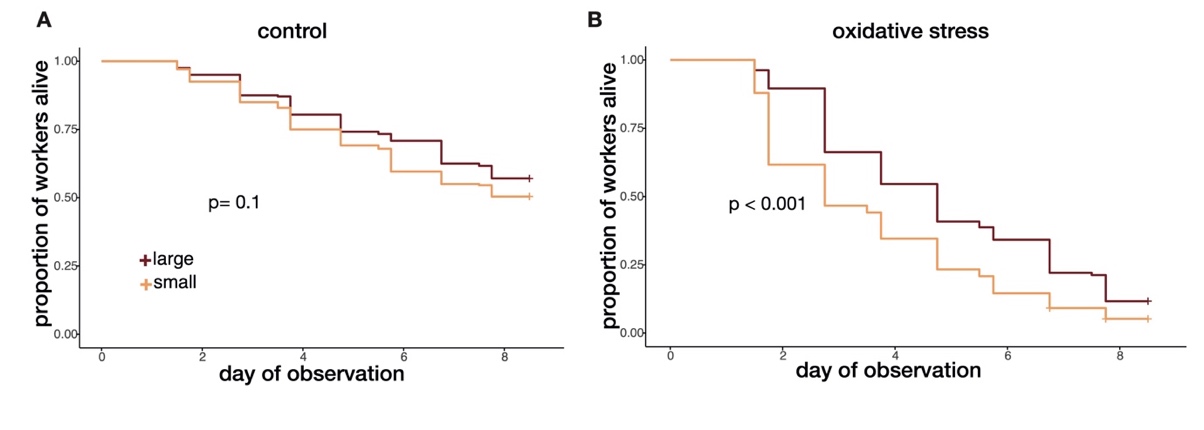

Figure S 2: Worker survival dependent to worker size within the oxidative stress experiment. A) Large (dark red line) and small (orange line) workers within the control treatment survived similarly long (X^2^= 2.69, p= 0.1). B) Large workers did survive better than small workers when subjected to paraquat- induced oxidative stress (X^2^= 46.11, p< 0.001).
