## Supplementary Figure S3 for "Queen number shapes worker longevity, fecundity and gene expression in the invasive, highly polygynous ant *Tapinoma magnum*"

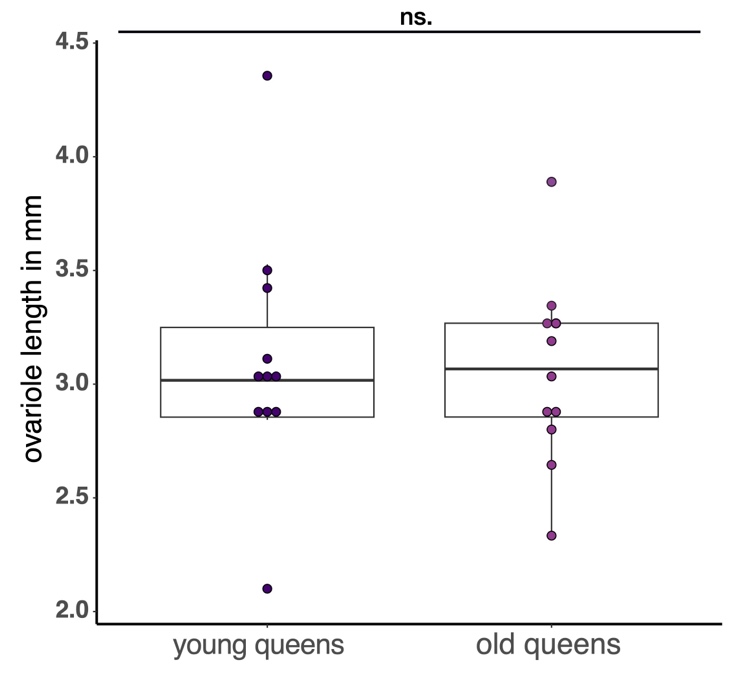

Figure S 3: Mean ovariole length of young and old Tapinoma magnum queens. Young and old queens had similar long ovarioles (LMER: X^2^= 0.02, p= 0.811).
