## Supplementary Material for "Queen number shapes worker longevity, fecundity and gene expression in the invasive, highly polygynous ant *Tapinoma magnum*"

***RNA extraction and Sequencing***

Large inside and outside workers from each experimental box were sampled after the 58-day survival experiments. The workers were frozen at -80°C between 12:00 and 16:00 and their fat bodies were dissected. We took the fat body from four individual workers and pooled their fat bodies to create 72 individual RNA pools. Each pool of four fat bodies was stored in 100µl of TRIzol™ LS Reagent (Invitrogen™) at -80°C until extraction. RNA was extracted in a 4:1 ratio of TRIzol: Chloroform: Isoamylalcohol followed by purification steps using the Qiagen RNA-easy Mini Kit. At the Beijing Genomics Institute (BGI), the High Sensitivity RNA Analysis Kit (Fragment Analyser) was used to check the quality of the extracted RNA. The libraries were prepared and sequenced by BGI using the Illumina HiseqXTen sequencing platform, yielding 150 bp paired- end reads with a sequencing depth of 23.36 ± 3 (mean ± sd) million reads.

For the queen dataset, we sampled and dissected six young and six old queens (12 total queens in total) and transferred brain and abdominal fat body tissues into 100µl of TRIzol™ LS Reagent (N= 12 brain samples; N=12 fat body samples). RNA extractions were conducted as described above and extracted RNA was checked using the Agilent 2100 Bioanalyzer, RNA 6000 Nano Kit at BGI. Library preparation and sequencing were similarly performed by BGI using the Illumina HiseqXTen sequencing platform, yielding in 150 bp paired- end reads with a sequencing depth of 23.58 ± 2.54 (mean ± sd) million reads.
