## Supplementary Table S1 for "Queen number shapes worker longevity, fecundity and gene expression in the invasive, highly polygynous ant *Tapinoma magnum*"

Table S 1: Significantly enriched GO Terms (molecular function) in the list of genes that were overexpressed in inside workers compared to outside workers including the respective p-values obtained from the Fisher’s exact test.

| **GO-ID** | **Term** | **Annotated** | **Significant** | **Expected** | **Fisher’s exact test p-value** |
| --- | --- | --- | --- | --- | --- |
| **GO:0016705** | oxidoreductase activity, acting on paired donors, with incorporation or reduction of molecular oxygen | 368 | 29 | 6.79 | 2.1E-09 |
| **GO:0005506** | iron ion binding | 377 | 25 | 6.96 | 3.2E-08 |
| **GO:0004497** | monooxygenase activity | 323 | 21 | 5.96 | 4.8E-07 |
| **GO:0005515** | protein binding | 4385 | 128 | 80.93 | 5.1E-07 |
| **GO:0003723** | RNA binding | 1254 | 49 | 23.14 | 9.9E-06 |
| **GO:0005509** | calcium ion binding | 496 | 22 | 9.15 | 0.00016 |
| **GO:0020037** | heme binding | 437 | 20 | 8.07 | 0.00021 |
| **GO:0008017** | microtubule binding | 131 | 9 | 2.42 | 0.00075 |
| **GO:0000981** | DNA-binding transcription factor activity, RNA polymerase II-specific | 179 | 8 | 3.3 | 0.03658 |
| **GO:0005085** | guanyl-nucleotide exchange factor activity | 139 | 6 | 2.57 | 0.04448 |
