## Supplementary Table S1 for "Queen number shapes worker longevity, fecundity and gene expression in the invasive, highly polygynous ant *Tapinoma magnum*"

Table S 2: Significantly enriched GO Terms (molecular function) in the list of genes that were overexpressed in outside workers compared to inside workers including the respective p-values obtained from the Fisher’s exact test.

| **GO.ID** | **Term** | **Annotated** | **Significant** | **Expected** | **Fisher’s exact test p-value** |
| --- | --- | --- | --- | --- | --- |
| **GO:0005515** | protein binding | 4385 | 209 | 123.95 | 1.3E-19 |
| **GO:0020037** | heme binding | 437 | 37 | 12.35 | 3.7E-09 |
| **GO:0004497** | monooxygenase activity | 323 | 31 | 9.13 | 4.3E-09 |
| **GO:0005506** | iron ion binding | 377 | 33 | 10.66 | 4.5E-09 |
| **GO:0016705** | oxidoreductase activity, acting on paired donors, with incorporation or reduction of molecular oxygen | 368 | 32 | 10.4 | 9.2E-09 |
| **GO:0005509** | calcium ion binding | 496 | 30 | 14.02 | 9E-05 |
| **GO:0004181** | metallocarboxypeptidase activity | 36 | 6 | 1.02 | 0.00047 |
| **GO:0004842** | ubiquitin-protein transferase activity | 170 | 13 | 4.81 | 0.00122 |
| **GO:0004252** | serine-type endopeptidase activity | 314 | 19 | 8.88 | 0.00166 |
| **GO:0008270** | zinc ion binding | 975 | 44 | 27.56 | 0.00167 |
| **GO:0005085** | guanyl-nucleotide exchange factor activi... | 139 | 10 | 3.93 | 0.00624 |
| **GO:0004674** | protein serine/threonine kinase activity | 158 | 11 | 4.47 | 0.01036 |
| **GO:0004222** | metalloendopeptidase activity | 207 | 12 | 5.85 | 0.01514 |
| **GO:0031267** | small GTPase binding | 72 | 6 | 2.04 | 0.01614 |
| **GO:0140359** | ABC-type transporter activity | 145 | 10 | 4.1 | 0.01809 |
| **GO:0043169** | cation binding | 2690 | 136 | 76.04 | 0.01945 |
| **GO:0016747** | acyltransferase activity, transferring g... | 446 | 15 | 12.61 | 0.02366 |
| **GO:0016298** | lipase activity | 111 | 9 | 3.14 | 0.03566 |
| **GO:0050660** | flavin adenine dinucleotide binding | 311 | 13 | 8.79 | 0.04072 |
